# Splicing-related Retinitis Pigmentosa mutations mimicked in *C. elegans* allow the identification of disease modifiers and drug screens

**DOI:** 10.1101/752956

**Authors:** Dmytro Kukhtar, Karinna Rubio-Peña, Xènia Serrat, Julián Cerón

## Abstract

CRISPR and the high conservation of the spliceosome components facilitate the mimicking of human pathological mutations in splicing factors of model organisms. The degenerative retinal disease Retinitis Pigmentosa (RP) is caused by mutations in distinct types of genes, including missense mutations in splicing factors that provoke RP in an autosomal dominant form (s-adRP). Using CRISPR in *C. elegans*, we generated mutant strains to mimic RP mutations reported in PRPF8 and SNRNP200. Whereas these inherited mutations are present in heterozygosis in patients, *C. elegans* allows the maintenance of these mutations in homozygosis, which is advantageous for genetic and drug screens. We found that *snrp-200*(*cer23*[V676L]) and *prp-8*(*cer14*[H2302del]) display pleiotropic phenotypes, including a reduced fertility. However, *snrp-200*(*cer24*[S1080L]) and *prp-8*(*cer22*[R2303G]) are weak alleles suitable for RNAi screens to identify genetic interactions, which would uncover potential disease modifiers. We screened a collection of RNAi clones for splicing-related genes and identified three splicing factors, *isy-1*/ISY1, *cyn-15*/PPWD1 and *mog-2*/SNRPA1 whose partial inactivation may modify the course of the disease. Interestingly, these three genes were acting as modifiers of *prp-8(cer22)* but no *snrp-200(cer24)*.

Finally, the strong allele *prp-8(cer14)* was used in a screen with FDA-approved drugs to find molecules capable of alleviating the phenotype. Instead, we detected drugs, as Dequalinium Chloride, which exacerbated the phenotype and therefore are potentially harmful for s-adRP patients since they may accelerate the progression of the disease.

## INTRODUCTION

Retinitis Pigmentosa (RP) is a rare disease with a prevalence of approximately 1 in 4000 individuals. Thus, this disease is causing progressive loss of vision to more than 1 million people worldwide. RP causes defects in photoreceptors (rods and cones) leading to apoptosis of these cells and degeneration of the retina (1). Although sporadic RP may occur, most of the RP patients present inherited mutations through diverse modes of genetic transmission. For all these RP familial cases, autosomal dominant is the most common form of genetic inheritance. To date, mutations in 26 genes have been linked with autosomal dominant Retinitis Pigmentosa (adRP) (2, 3), and 7 of these genes encode splicing factors.

The spliceosome is probably the most complex cellular machine. More than 200 proteins are part of this highly dynamic macromolecular complex that removes introns from eukaryotic pre-mRNAs (4). The spliceosome consists of small nuclear ribonucleoproteins (complexes of snRNAs (U1, U2, U4, U5 and U6) and proteins) that form the core of the spliceosome and have multiple associated proteins. Interestingly, the seven splicing-related genes linked with adRP, encode proteins present in the U4/U6·U5 tri-snRNP complex, which is involved in the assembly of the catalytic spliceosome (5). These seven proteins are PRPF3, PRPF4, PRPF6, PRPF8, PRPF31, SNRNP200 and RP9. All genes, but RP9, encode proteins highly conserved from *C. elegans* to humans.

RP patients carrying mutations in these splicing factors do not show any other pathologies or phenotypes associated producing a paradox that is not completely solved yet: mutations in genes expressed ubiquitously present a disease in a specific tissue, the retina (6). No reliable treatment or therapy is available to treat RP patients. Thus, a better understanding of the molecular and cellular consequences of s-adRP mutations would help to address this problem, in s-adRP patients at least. However, there are two main handicaps in the study of these mutations, (i) the time-scale in which the disease progresses, and (ii) the essential nature of splicing factors that often hampers genetic and cellular studies. In this study, we propose the use of CRISPR in *C. elegans* to investigate s-adRP mutations and to explore therapies for RP. Importantly, we have found that these mutations are viable in homozygosis, facilitating functional studies and large-scale screens.

RP patients within the same family, and therefore carrying the same mutations, often present the onset of the disease at different age and the disease progresses differently among the family members. Such variability can be explained by the existence of modifiers, which are mutations or polymorphisms in other genes that influence onset and progression of the disease (7). As an example, family members harboring the same mutation in PRPF31, present the ages of onset of the disease between 21 and 63 (8). Our study proposes the use of *C. elegans* as model to *(i)* identify potential modifiers of the splicing-related RP, which could contribute to a better prognosis of patients, and (ii) to identify drugs that could alleviate or accelerate the disease.

## RESULTS

### Splicing-related RP missense mutations engineered in *C. elegans*

PRPF8 and SNRNP200 are two of the seven genes associated to autosomal dominant Retinitis Pigmentosa (adRP) (9). Several mutations in these two genes have been identified in distinct families at different countries (10–16). Whereas PRPF8 mutations are in the *C-terminal* end of the protein, SNRNP200 mutations are present in central helicase domains (**Figure 1A**).

**Figure 1.**
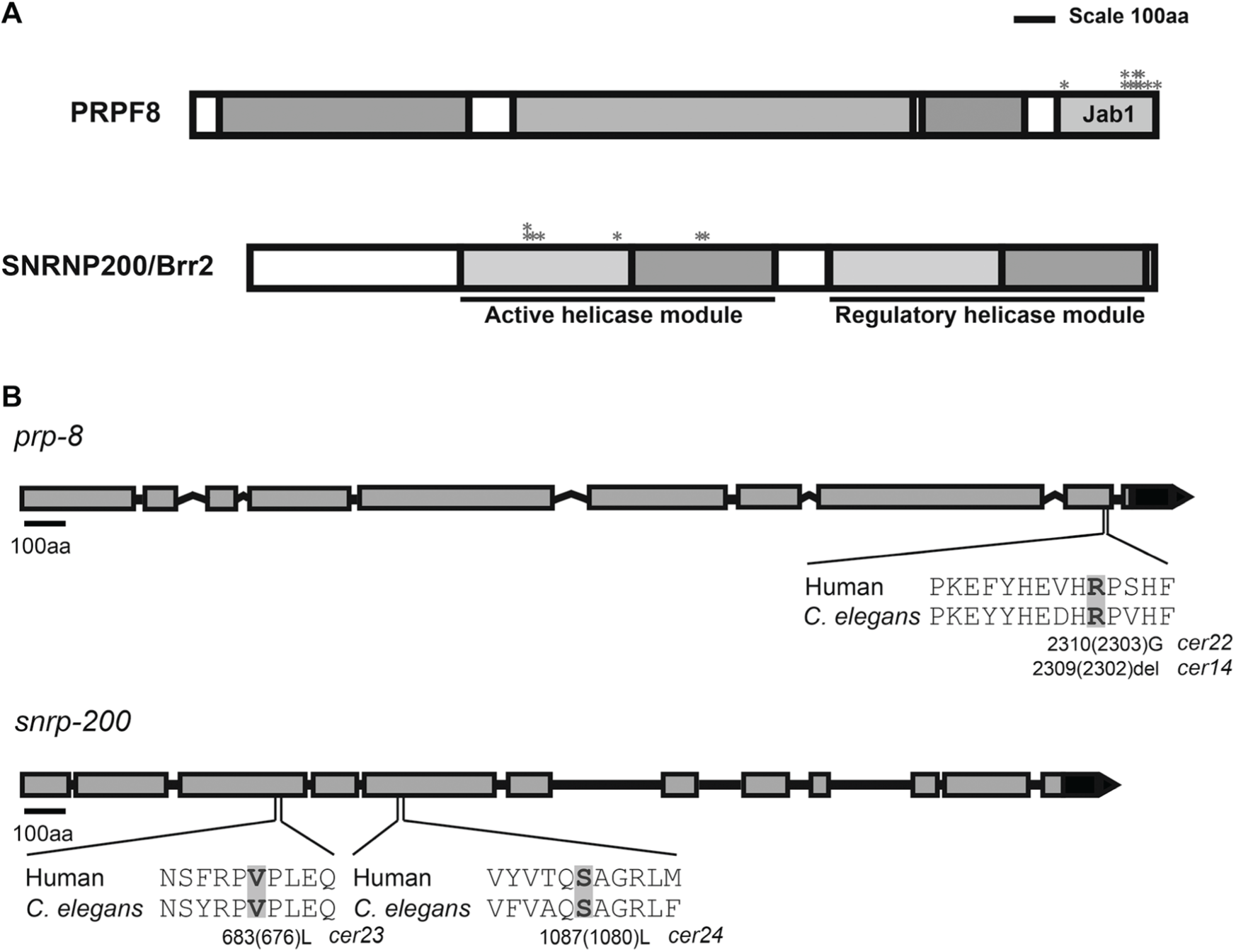
Domain structure and RP mutations in PRPF8 and SNRNP200/Brr2. (A) Scale representation of PRPF8 and SNRNP200/Brr2 human genes. The position of mutations linked to adRP is indicated with asterisks. Jab1 domain is located at the C-terminal end of PRPF8. Helicase domains are indicated in SNRNP200/Brr2. (B) Schematic representation of *prp-8* and *snrp-200 C. elegans* genes. Amino acids affected by the RP mutations generated in this study and their worm counterparts are shaded. *cerxx* indicates the alleles generated in this study.

We first produced the *C. elegans* missense mutation corresponding to PRPF8[R2310G], which is a mutation that has been identified in South African and Spanish families (17, 18) (**Figure S1A**). As a result of this attempt, we isolated the *C. elegans prp-8(cer22[R2303G])* allele that resemble the PRPF8[R2310G] mutation (**Figure 1B**). Unintentionally, we also identified the *prp-8(cer14[H2309del])* allele, which consist of a 4nt deletion/1nt insertion (or small indel) that remove the Histidine 2309 (2302 in *C. elegans)*, an amino acid that is important to maintain the interactions of PRPF8 with other proteins, and it is also mutated in s-adRP patients (17, 19) (**Figure 1B**) **(Figure S1)**.

The C-terminal end of PRPF8, in which adRP mutations are located, is the region that interacts with SNRNP200 (10). We decided to reproduce adRP mutations in the SNRNP200 *C. elegans* ortholog *snrp-200* reasoning that their molecular effect could be similar to PRPF8 mutations. Thus, we produced the *snrp-200* alleles *cer23*[V676L] and c*er24*[S1080L], corresponding to patient’s mutations SNRPN200[V683L] and SNRPN200[S1087L] respectively (**Figure 1B**) (**Figure S1B**).

### Phenotypes of adRP mutations in *C. elegans*

The four mutant strains mentioned above can be maintained in homozygosis. However, we observed that these strains seemed to grow slower. We formally tested this observation at distinct temperatures, and we found that the four strains presented a slight postembryonic developmental delay, more prominent at 25°C and in animals carrying the *prp-8(cer14)* and *snrp-200(cer23)* alleles (**Figure 2A, S2**) (**Table 1**). These two alleles, *cer14* and *cer23* also displayed a variety of developmental defects at low penetrance, including larval arrest, larval morphological defects and dumpy animals (**Figure S3**). We also scored the fertility of the s-adRP mutants at 25°C since mutations in splicing factors commonly display stronger penetrance at that temperature (20, 21). We observed that *prp-8(cer14)* and *snrp-200(cer23)* animals presented a reduction of the progeny size (**Figure 3A, 3B**) (**Table 1**). Then we studied the fertility of *prp-8(cer14)* at permissive (15°C) and restrictive (25°C) temperatures and concluded that *cer14* is a temperature-sensitive allele that cause sterility at 25°C and a reduced brood size at 15°C (**Figure 3B**). We investigated the cause of the sterility of *prp-8(cer14)* mutants and found that these animals do not develop the germline properly, presenting shortened germline that does not even made the U-turn (**Figure 3C**).

**Table 1.** Main phenotypes of *prp-8* and *snrp-200* s-adRP mutations generated by CRISPR. Phenotype abbreviations: Ste: sterility, Rbs: reduced brood size, Gro: developmental delay.

| Type of mutation | Allele | Description | Phenotype |
| --- | --- | --- | --- |
| ***prp-8*** |  |  |  |
| point mutation | *cer22* | R2310G (R2303G) | Gro |
| indel | *cer14* | H2309(H2302) aa loss | Rbs, Ste, Gro |
| ***snrp-200*** |  |  |  |
| point mutation | *cer23* | V683L(V676L) | Rbs, Gro |
| point mutation | *cer24* | S1087L(S1080L) | Gro |

**Figure 2.**
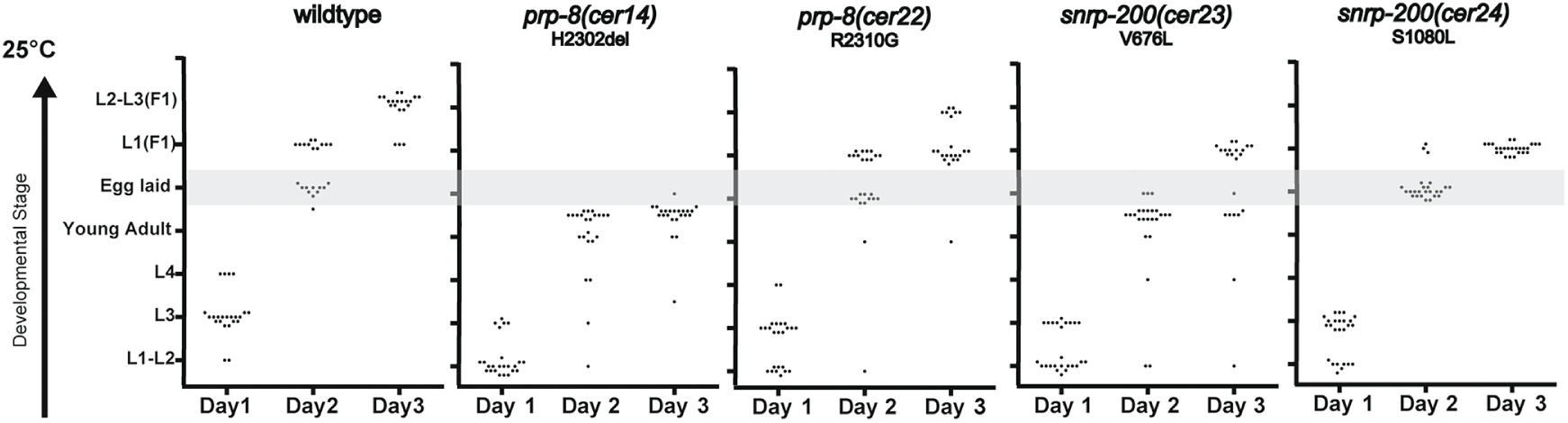
RP mutants are viable but present developmental delay. *prp-8* and *snrp-200* mutations provoked delay in postembryonic development. Developmental progress of singled, mutant and wild-type, L1 larvae (n=25) was monitored every 24 h for 3 days at 25°C. Every dot represents a worm. Shaded area helps to visualize the presence or absence of embryos in the plate.

**Figure 3.**
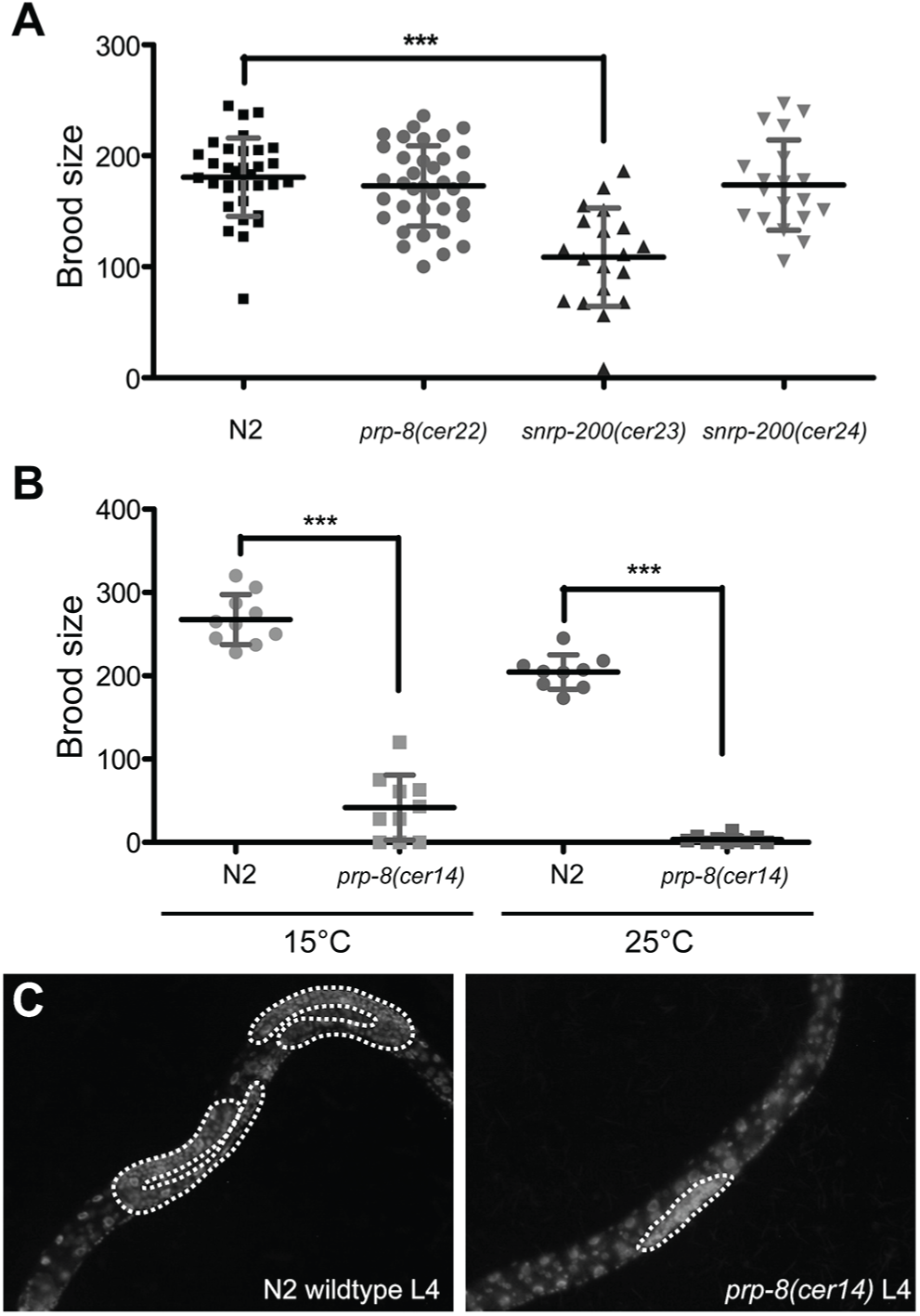
RP mutations have different effects in *C. elegans* fertility. (A) Brood size of RP mutants at 25°C. Worms were grown and singled out at L4 stage at the corresponding temperature (n≥20). Adult animals were transferred to new plates every 8-12 hours and the number of larvae (F_1_) determined. Brood size of *snrp-200(cer23)* animals is significantly reduced compared with wild-type (p value<0.0001; unpaired t test). Error bars indicate the standard deviation. (B) Progeny of *prp-8(cer14)* animals. Worms were grown at 15°C or 25°C until reaching the L4 stage, singled out and kept at their corresponding temperature (n≥10). Adults were transferred to new plates every 8-12 hours and the number of larvae (F_1_) determined. Brood size of *prp-8* mutants is reduced at both temperatures (p value<0.0001; unpaired t test). Error bars indicate the standard deviation. (C) Representative images of DAPI staining of WT and *prp-8(cer14)* L4 animals. Discontinued white line marks the germline, which is smaller in *prp-8(cer14)* mutants.

As a summary of the phenotypic characterization, we described two types of mutations, being *prp-8(cer22)* and *snrp-200(cer24)* weak alleles, and *prp-8(cer14)* and *snrp-200(cer23)* strong temperature-sensitive alleles. Weak alleles are useful to screen for synthetic interactions as means of identification of modifiers of the disease whereas the strong alleles are convenient in the search of drugs that can alleviate the mutant phenotype.

### RNAi screen for modifiers of s-adRP mutations

Since s-adRP mutations affect core proteins of the spliceosome, we reasoned that reduced or altered functions of other spliceosome components might increase the harmful effect of s-adRP mutations. Consequently, partial loss-of-function mutations or polymorphisms in other splicing factors could act as modifiers of the disease. Thus, we made a library of RNAi clones targeting 98 splicing factors that were assayed in wild-type, and in *prp-8(cer22)* and *snrp-200(cer24)* animals, which are the two mutant strains harboring weak alleles. All the RNAi clones were tested in duplicate on solid media, in 24-well plates, in wild-type and mutant backgrounds. In this first screen, we identified three genetic interactions with the *prp-8(cer22)* mutants (**Figure 4A**). No interactions were observed with the *snrp-200(cer24)* allele.

**Figure 4.**
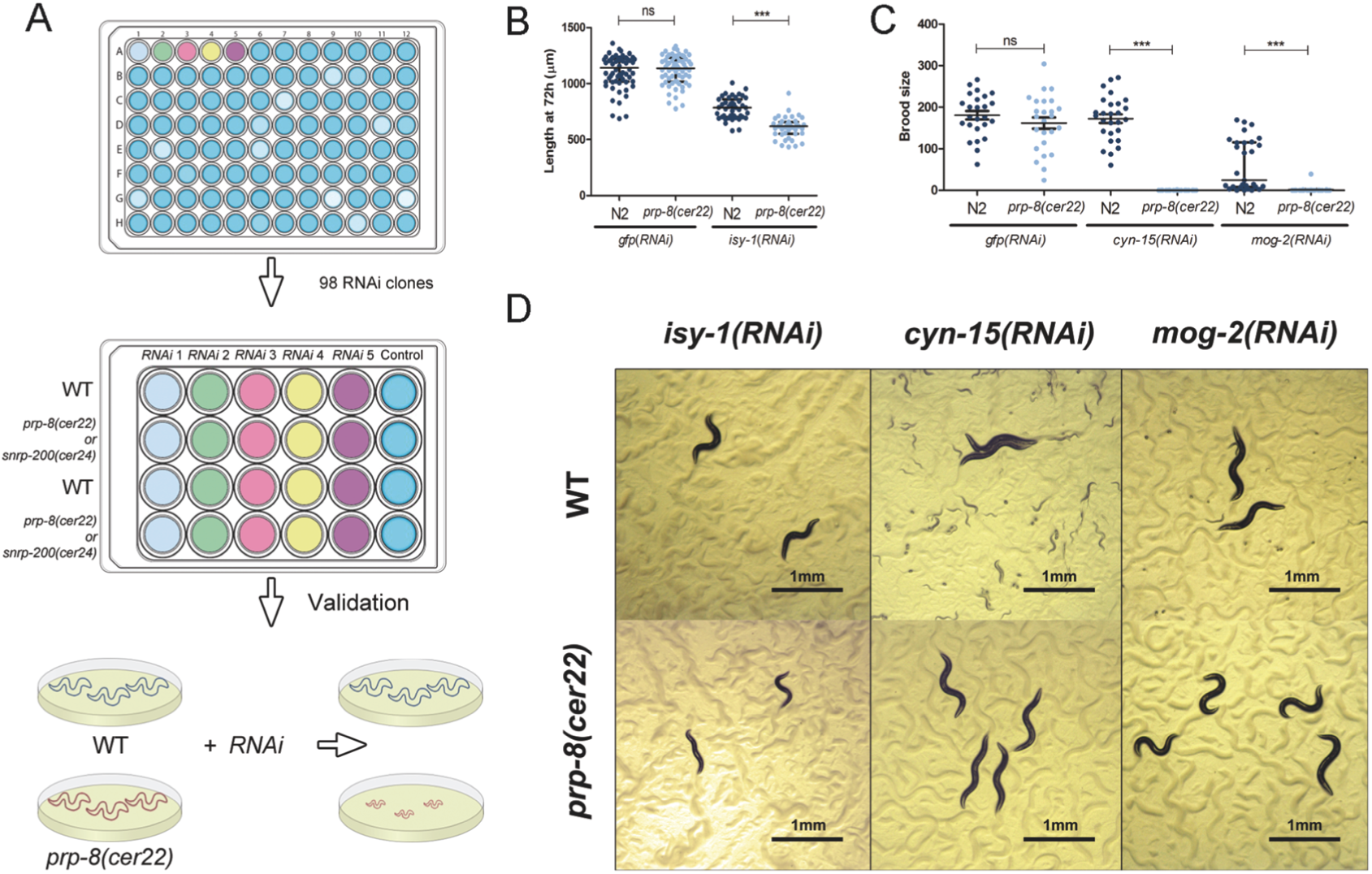
RNAi screen of splicing-related genes identifies genetic interactions with *prp-8(cer22*). (A) Schematic representation of RNAi screen procedure. We assayed 98 splicing-related genes on wild-type (WT), and *prp-8(cer22)* and *snrp-200(cer24)* mutants, in 24-well plate format. *isy-1, cyn-15* and *mog-2* were identified as enhancers of the phenotype in *prp-8(cer22*) mutants, and were validated in individual agar plates. (B) Length of WT and *prp-8(cer22)* animals at 72 h after hatching, grown at 25°C (n≥61, N=3). Each dot represents the body length of a single worm, bars represent median with interquartile range (IQR). (C) Progeny of WT and *prp-8(cer22)* mutants (n≥25, N=2). Each dot represents the offspring of a single worm, bars represent median with IQR. (D) Representative images of WT and *prp-8(cer22)* animals fed with RNAi clones for *isy-1, cyn-15* and *mog-2*. Kruskal-Wallis with Dunn’s post-hoc analysis: *ns* indicates not significant and *\*\*\** p≤0.001.

We validated and quantified these genetic interactions in bigger plates and in duplicates. We confirmed that *prp-8(cer22)* animals were smaller (developmental arrest) than wild type when exposed to *isy-1(RNAi)* (**Figure 4B, 4D)**. Moreover, we determined that *prp-8(cer22)* mutants presented a reduced fertility in *cyn-15(RNAi)* and *mog-2(RNAi)* when compared with control animals (**Figure 4C, 4D)**.

In summary, we identified genetic interactions that were specific for a mutant strain and found that these interactions affect different aspects of worm development since one produce growth delay, and the other two reduced fertility.

Next, we wondered if common polymorphisms of ISY1 (*isy-1* human ortholog) and PPWD1 (*cyn-15* human ortholog) could functionally interact with the *prp-8*(*cer22*[R2303G]) mutation (22). We selected residues with polymorphisms from the top 20 most prevalent polymorphisms in healthy humans using *in silico* tools that predict the effect of amino acid changes on proteins (23, 24). By CRISPR, we introduced the single amino acid changes *isy-1(cer114[G170S])* and *cyn-15(cer119[D74Q])* and these mutations were combined with *prp-8(cer22)* to produce double mutant strains (**Figure S4**). We did not observe any synthetic interaction, but our experiment illustrates the possibility of easily testing the effect of human polymorphisms, which may potentially modify the onset or the progression of the disease.

### Drugs screen

RP patients present a progressive loss of visual field through their life and we reasoned that a drug with an even slight positive effect on our *C. elegans* mutants would cause delay in the progression of the disease, resulting in a better quality of life for patients. We used a library with 929 drugs, 91.8% approved by the FDA, and therefore hits that would be validated in other pre-clinical models would easily be transferred to the clinic. Synchronized L1 animals were exposed to 10 µM of each drug and grown on dead bacteria at 25°C in liquid medium. The effect of the drug in wild-type and *prp-8(cer14)* mutant animals was scored at adult stages (**Figure 4A**). To better characterize the efficacy of our collection of drugs, after three days of exposure we quantified the locomotor activity of the worms for 15 minutes using the wmicrotracker device (25). We observed that drugs causing an obvious phenotype also affected the locomotor activity of the animals and, consequently, drug effect could also be detected by an automated detection of locomotion (**Figure S5**).

The aim of our screen was to find any molecule capable of rescuing the sterile phenotype of *prp-8(cer14)*. We observed a phenotype in wild-type animals for 146 of the 929 drugs tested (about 16%), validating the functional activity of the library, but we did not observe any clear rescue of the *prp-8(cer14)* sterility.

Instead, we observed that four molecules (Doxycycline hydrochloride, Flutamide, Dequalinium Chloride and Dronedarone hydrochloride) caused a stronger effect on mutants than in the wild-type animals. Such finding is also of clinical interest since the use of these drugs in humans is approved and RP patients could use a drug that may accelerate the progression of their disease. We further tested these four drugs on solid media with all s-adRP strains and only one of these drugs, Dequalinium Chloride (DQ), which is commonly used as antiseptic, was consistently harmful to s-adRP mutants. After validations, the adverse effect of DQ was consistently reproduced in the two *snrp-200* mutants only, and at specific concentrations of the drug (**Figure 5B, 5C, S6**).

**Figure 5:**
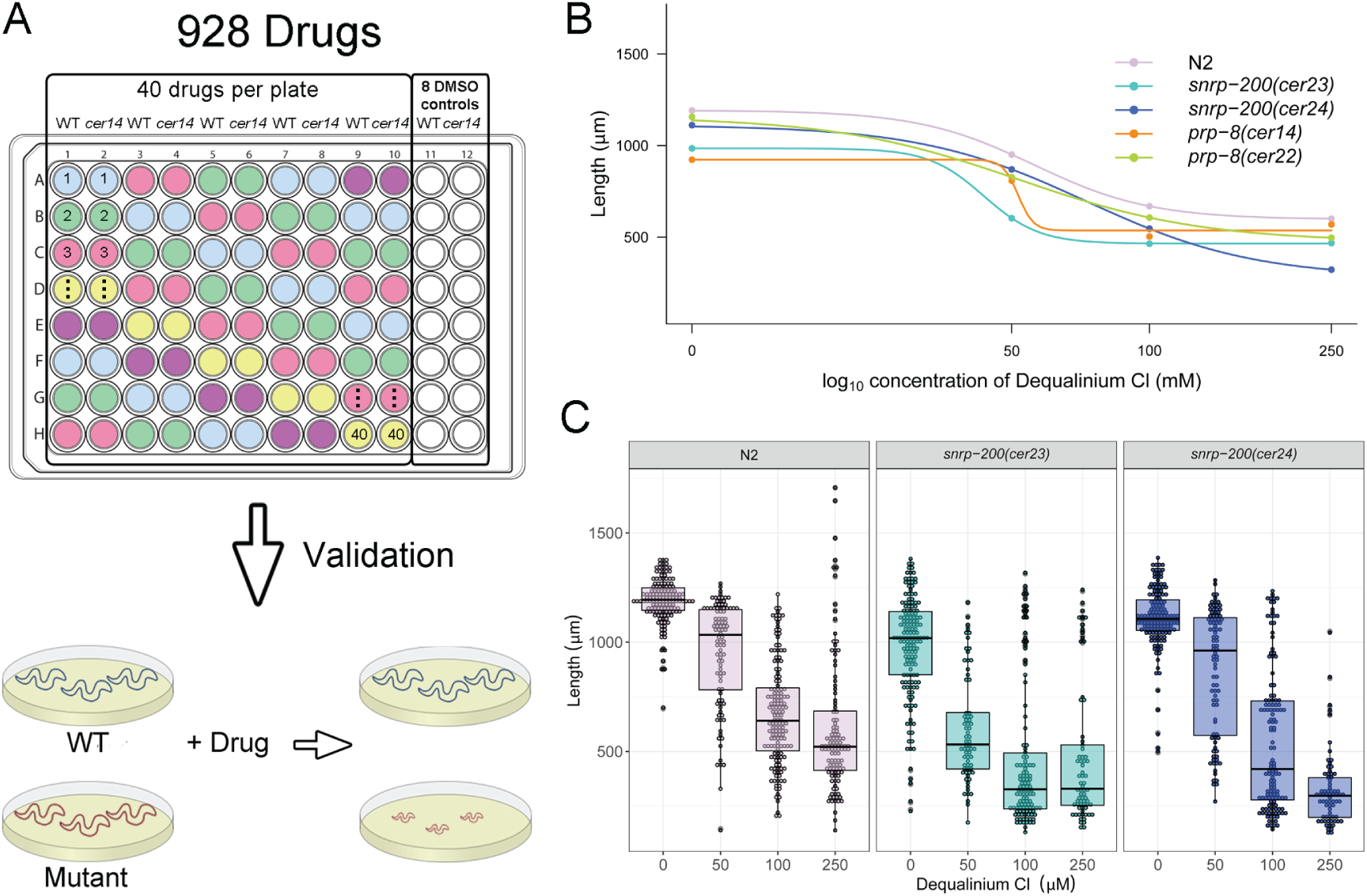
Screen of FDA-approved and epigenetic drugs. (A) Schematic representation of the drug screen procedure. Drugs were tested in duplicates, in liquid medium with 0,5% DMSO, at a rate of 40 drugs per each 96-well plate. Hits were validated on individual agar plates. (B) Dose-response curve of Dequalinium Chloride (DQ) in *C. elegans* wild-type (N2) and s-adRP strains. Each dot represents the median of the length at each condition (n≥69) and the line Log-logistic distribution (N=3, except N=2 at 50 µM). (C) Worm length of wild-type (N2) and *snrp-200* mutant strains upon increasing DQ concentrations. Each dot represents the length of an individual worm (n≥69). Box-plot indicates the median with the IQR and whiskers ± 1.5 product of IQR (N=3, except N=2 at 50 µM).

## DISCUSSION

There are six splicing factors mutated in RP that are highly conserved in *C. elegans*. We chose PRPF8 and SNRNP200 because these two proteins physically interact through the C-terminal region of PRPF8, where the s-adRP mutations are located (10). Moreover, RNAi of their orthologs in *C. elegans*, along with PRPF6, showed the most penetrant phenotypes (26). These three genes encode proteins of the U5 complex and, based on our previous RNAi experiments, would be more prone to harbor missense mutations with a strong phenotype than the other three splicing factors (PRP3, PRPF31 and PRPF4), which are part of the U4 complex. In fact, PRPF31 is the only s-adRP gene that admits a deletion of the gene causing RP by haploinsufficiency (27).

In humans, s-adRP mutations cause defects that provoke degeneration of photoreceptors in the retina through a very slow process (28). Such slowness is a handicap in the modeling of the disease and hampers the understanding of the molecular mechanisms of the pathology.

As drawback to our animal model, *C. elegans* lacks photoreceptor-like cells. Photoreceptors are non-dividing cells that are constantly renewing components of the outer segment membrane disc, which implies an intense transcriptional activity and metabolic rate (29). Similarly, cells in *C. elegans* also present increased metabolic rate (30) and high transcriptional levels during larval development (31). Importantly, since *C. elegans* s-adRP mutations can be maintained as homozygous, it is possible to perform experiments with hundreds or thousands of mutant animals allowing the screening for genetic modifiers or drugs that could either alleviate or exacerbate the phenotype.

Our *C. elegans* models for s-adRP mutations rely on a single change on amino acids that are conserved from nematodes to human. Such extraordinary conservation would imply maintenance of the function and therefore the missense mutations could have similar consequences at the molecular level in humans and in nematodes. *C. elegans* has already been exploited as model for s-adRP (26) and would be further exploited in the future if functional replacement of *C. elegans* s-adRP genes at its endogenous locus can be achieved with their human counterparts. The challenge here would be to express a functional human splicing factor at endogenous levels without causing any phenotype in the nematode. Alternatively, if the functional replacement cannot be complete, a fraction of the protein could be replaced. Thus, we recently humanized the pladienolide binding domain of *sftb-1*/SF3B1 to make *C. elegans* sensitive to this drug without causing any other effect in the animal (32).

The existence of modifiers of s-adRP would explain the distinct penetrance of the disease. A potential modifier has been identified for s-adRP patients with mutations in PRPF31 (33). Asymptomatic patients with PRPF31 mutations correlate with lower levels of CNOT3, which is a transcriptional regulator of PRPF31. In this case, CNOT3 is a modifier in *trans*-that would modify the effect of any mutation in the gene PRPF31, but not mutations in other s-adRP genes.

Invertebrate animal models as *C. elegans* and *Drosophila melanogaster* are powerful genetic systems to study genetic modifiers. In the case of RP, *Drosophila* has already been used to identify candidate genetic modifiers of a missense mutation in the fly ortholog of human rhodopsin (34).

Our RNAi screen identified genetic interactions with *prp-8(cer22)* but we did not obtain any hit in the *snrp-200(cer24*) background. This result reinforces the idea that genetic modifiers of the s-adRP disease could be gene specific, or even mutation-specific since diverse PRPF8 s-adRP mutations have distinct effect in PRPF8 stability and location (35). Still, the existence of universal modifiers of s-adRP is possible but they have not been found yet. Therefore, the lack of a list for global modifiers of the disease complicates the prognosis of RP patients.

RNAi inactivation, which mimic a partial loss-of-function effect, uncovered a synthetic interaction between *isy-1, cyn-15* and *mog-2* and the *prp-8(cer22)* allele but did not show any synthetic effect in *snrp-200(cer24)* mutant animals. Interestingly, the genetic interaction of *prp-8(cer22)* with *isy-1*/ISY1 gave a different phenotype (growth defects) than the interactions with *mog-2*/SNRPA1 and *cyn-15* /PPWD1 (reduced fertility). Since PRPF8 is the biggest protein of the spliceosome, and it is present in most of the splicing steps (36), the activity of the candidate genetic modifiers may not be restricted to a single splicing mechanism. ISY1 is part of the NTC complex that participates in precatalytic and catalytic spliceosome. SNRPA1 is at the U2 complex, whose main role is the recognition of the 3’ splice site, but it is also present in the catalytic spliceosome (C complex). PPWD1 is one of the Peptydil Prolyl isomerases of the spliceosome and it is also present in the catalytic spliceosome (37, 38).

We mimicked in *C. elegans* two human polymorphisms that may affect the structure of ISY1 and PPWD1, but we did not detect any interaction with *prp-8(cer22)*. However, any missense mutation found in the conserved amino acids of ISY1, PPWD1 and SNRPA1, which are not prompted to present polymorphisms (**Figure S4**), would be considered as a candidate for being a modifier of s-adRP caused by PRPF8[R2303G].

*C. elegans* is a well-established model for drug discovery (39, 40). Many drugs with an impact on human cells also cause an effect in *Drosophila* and *C. elegans* (41, 42). We assayed a drug library with about 1000 small molecules, most of them FDA-approved, and 16% of them caused an obvious phenotype in *C elegans*. Our initial objective was to identify a drug capable of rescuing the temperature sensitive sterility of *prp-8(cer14)* mutants. However, we obtained an unexpected result since drugs produced a stronger phenotype in mutant animals than in wild type. Such result is also clinically relevant since these drugs could have a harmful effect in certain patients. We further validate such negative effect for Dequalinium (DQ), a cytotoxic drug used as antiseptic and disinfectant (43). DQ and its derivates has been studied for the treatment of distinct cancer types (44, 45), the use of nanosomes to induce apoptosis by production of ROS (46), and the mitochondrial targeting in nanomedicine (47). Similar to genetic modifiers, the negative effect of some drugs would be gene and/or mutation specific, but our result certainly opens a new venue to study the potential damaging effect of common drugs that may accelerate the progression of the disease in some patients.

## MATERIAL AND METHODS

### *C. elegans* strains and maintenance

Worms were cultured and managed following standard methods (48). Thus, worms were grown on NGM (Nematode Growth Media) agar plates previously seeded with an overgrown culture of the *Escherichia coli* strain OP50 at temperatures between 15 and 25°C depending on the experiment. Synchronization of worms was done following the sodium hypochlorite treatment (49).

Bristol N2 was used as the wild type strain. All other strains are listed next: CER255 (*prp-8(cer14*[2303del]*) III*), CER240 (*prp-8(cer22*[R2303G]*) III*), CER256 *snrp-200(cer23*[V676L]*) II*), CER248 (*snrp-200(cer24*[S1080L]*) II*), CER456 (*isy-1(cer115*[G170S]*) V*), CER465 (*cyn-15(cer119*[D74Q]*) I*), CER544 (*prp-8(cer22*[R2303G]*) III; isy-1(cer115*[G170S]*) V*) and CER545 (*prp-8(cer22*[R2303G]*) III; cyn-15 (cer119*[D74Q]*) I*).

### Brood size and DAPI staining

For brood size assessment L4 larvae of each strain were singled out in 35-mm NGM agar plates and transferred to a fresh plate daily until egg laying stopped. Two days after P_0_ was removed the number of F_1_ larvae from each plate was manually scored. DAPI staining was performed recovering worms in M9 and placing them onto a slide. Next, residual M9 was removed and 90% ethanol was added and let for around two minutes moment where DAPI-Fluoromount-G^®^ (SounthernBiotech ref: 0100-20) was added and afterward a coverslip was placed on the top of the slide. Pictures were taken on a Nikon ECLIPSE TI-s inverted microscope attached to a Nikon DS-2Mv camera using NIS Elements 3.1, SP3 software.

### CRISPR editing

*prp-8* and *snrp-200* mutant worm lines were generated via CRISPR Cas9-triggered homologous recombination following previously described methods (51, 52). All reagents including crRNAs, Alt-R™ CRISPR-Cas9 tracrRNA, Cas9 Nuclease 3xNLS and primers were obtained from Integrated DNA Technologies (IDT). crRNAs containing a guide sequence with an adjacent protospacer motif (PAM) was injected with purified Cas9 enzyme and *dpy-10* crRNA with ssODN *dpy-10(cn64)* as a co-injection marker for each mutation.

To generate the *prp-8(cer22*[R2303G]) mutation the crRNA was located in the 9th exon of *prp-8* (AGTACTATCATGAAGATCAT<u>CGG</u>). Single point mutation was identified by single worm genomic PCR of *dpy-10* Rol mutants using specific primers (Fwd-TATGGTGTATCGCCACCTGA; Rev-GAAGTGAACCGGCCGATG for wild-type; Rev-GAAGTGAACCGGGCCGTG for the mutant), alternatively a *BceAI* restriction site was also located in the mutant strain. The small indel allele *cer14*, obtained with the same crRNA, is identified using the specific reverse primer TTATGGAAGTGAACCGGCCC.

The *snrp-200(cer23*[V676]) mutant was created through homology-directed repair using a crRNA guide sequence where the PAM was within the *snrp-200* exon 3 (TCCAATATATTGTTGCTCGA<u>GGG</u>). The mutant was identified using specific primers (Fwd-GTGATCGTTTGTACGCCGGA, Rev-TATATTGTTGCTCGAGGGGAAC for wild-type; Rev-TACTGCTGTTCCAGAGGGAG for the mutant). Allele *cer24*[S1080] was generated using a crRNA guide sequence located in the 5^th^ exon of *snrp-200* (GTCTTCGTGGCTCAGAGTGC<u>CGG</u>). Identification of the mutant was made by simple PCR using specific primers (Fwd-CTCATCACAAATCACTCGGAG; Rev-GAAGAGTCGTCCGGCACTC for wild-type; Rev-GAAGAGTCGTCCGGCGAGT).

*isy-1(cer115*[G170S]) and *cyn-15(cer119*[D74Q]) were generated as described above. crRNA guide sequences used for injection: *isy-1* GGTTACTTGGATGACGAAGATGG and *cyn-15* ATAATCACTGCAAGCGTCGATGG. The identification of mutant lines was done by PCR with specific primers: *isy-1* (common Rev-CCGCTCAGTTTACTTGTATT; Fwd-TTACTTGGATGACGAAGATGG for wild-type; Fwd-GTTACCTTGACGATGAGGACTC for mutant) and *cyn-15* (common Rev-CCTCAATTATCTTGACTGGCG; Fwd-TAATCACTGCAAGCGTCGAT for wild-type; Fwd-TAATCACAGCGTCTGTGCAA for mutant).

### Developmental delay assay

Worms were synchronized using the sodium hypochlorite treatment (49). Each strain was seeded in an OP50 plate and let recover for 1 h at 20°C and 25°C. After that, they were singled onto small plates containing OP50 bacteria and grown at the corresponding temperature.

The stage of the worms was determined every 24 h during a 3 days period. Stage determination was based on size and development of body structures like vulva and the size of the gonad.

### RNAi screen

128 splicing-related RNAi clones were selected from the literature (21). Bacteria expressing dsRNA was obtained from ORFeome library (53) or Ahringer library (54), and it was authenticated assessing the size of the insert by PCR, moreover 6 randomly selected clones were Sanger sequenced prior to its usage, resulting in a total of 98 validated clones that were used in RNAi by feeding assays. The screen was done in 24 well plates containing NGM supplemented with Tetracycline 12,5 µg/ml, Ampicillin 50 µg/ml and 3 mM IPTG at 25°C (RNAi plates). Each clone was tested in duplicates with 10 to 20 of either WT or mutant (*prp-8(cer22) or snrp-200(cer24)*) worms per well, from L1 synchronized stage. *gfp*(RNAi) was used as negative control. The scoring of the worms was done at 72 h and 96 h post seeding.

The validation of the interaction with *isy-1(RNAi)* was done at 25°C in 55-mm RNAi plates. 72 h post-seeding 35x magnified pictures of the plate were taken using stereoscopic microscope NIKON SMZ800 attached to a DS-2MV camera and were measured manually by drawing the central line of the worm in NIS Elements 3.10, SP3 software. *cyn-15(RNAi)* and *mog-2(RNAi)* brood size was assessed as previously described using RNAi plates.

### Drug screen on liquid in 96-well Plates

A total of 929 drugs: 853 FDA approved and 76 epigenetic drugs, obtained from Selleck Chemicals at 10 mM (www.selleckchem.com), were tested. 96-wells plates containing 50 µM of each drug in S-basal supplemented with 5 µg/ml Cholesterol, 50 µg/ml Ampicillin, 12,5 µg/ml Tetracycline, and OP50 as food source were used. Around 10 either WT or *prp-8(cer14)* L1 synchronized worms were seeded to each well and kept in a humidified chamber at 25°C. The scoring was done at day 3 and day 4 by visual observation and 15 min wmicrotracker measurement. DMSO 0,5% was used as negative control. Potential candidates were retested under the same condition in triplicates.

### Drug validation in agar plates

Validation was done in duplicates in 35-mm plates containing NGM with OP50 that was freeze-thawed three times at −80°C as food source. Commercially available drugs from Sigma-Aldrich manufacturer were used: Dequalinium chloride (ref: PHR1300), Flutamide (ref: F9397), Dronedarone hydrochloride (ref: D9696) and Doxycycline hyclate (ref: D9891). Drugs were added on the top of the agar and kept at 4°C overnight to allow its diffusion. Around 50 L1 synchronized worms of each strain were added to each plate and kept at 25°C. At 48 h post-seeding pictures of the worms were taken using Zeiss Axiocam ERc 5s under Zeiss Stemi 305 stereomicroscope and measured manually by drawing the central line of the worm in Fiji ImageJ (v1.51w) software.

## ACKNOWLEDGMENTS

We acknowledge the members of the Cerón Laboratory for helpful discussions in lab meetings, and Nicholas Stroustrup, Michael Krieg and Sonia Guil, for their ideas and comments as members of DK PhD committee.

This work have been supported by a grant from the ISCIII to JC [grant reference PI15-00895] co-funded by FEDER funds/European Regional Development Fund (ERDF) — a way to build Europe, and thanks to finantial support from Fundación ONCE. We thank CERCA Program / Generalitat de Catalunya for their institutional support. DK has an FI AGAUR fellowship from Generalitat de Catalunya and XS has an FPU PhD fellowship from MINECO.

---

**Figure S1.**
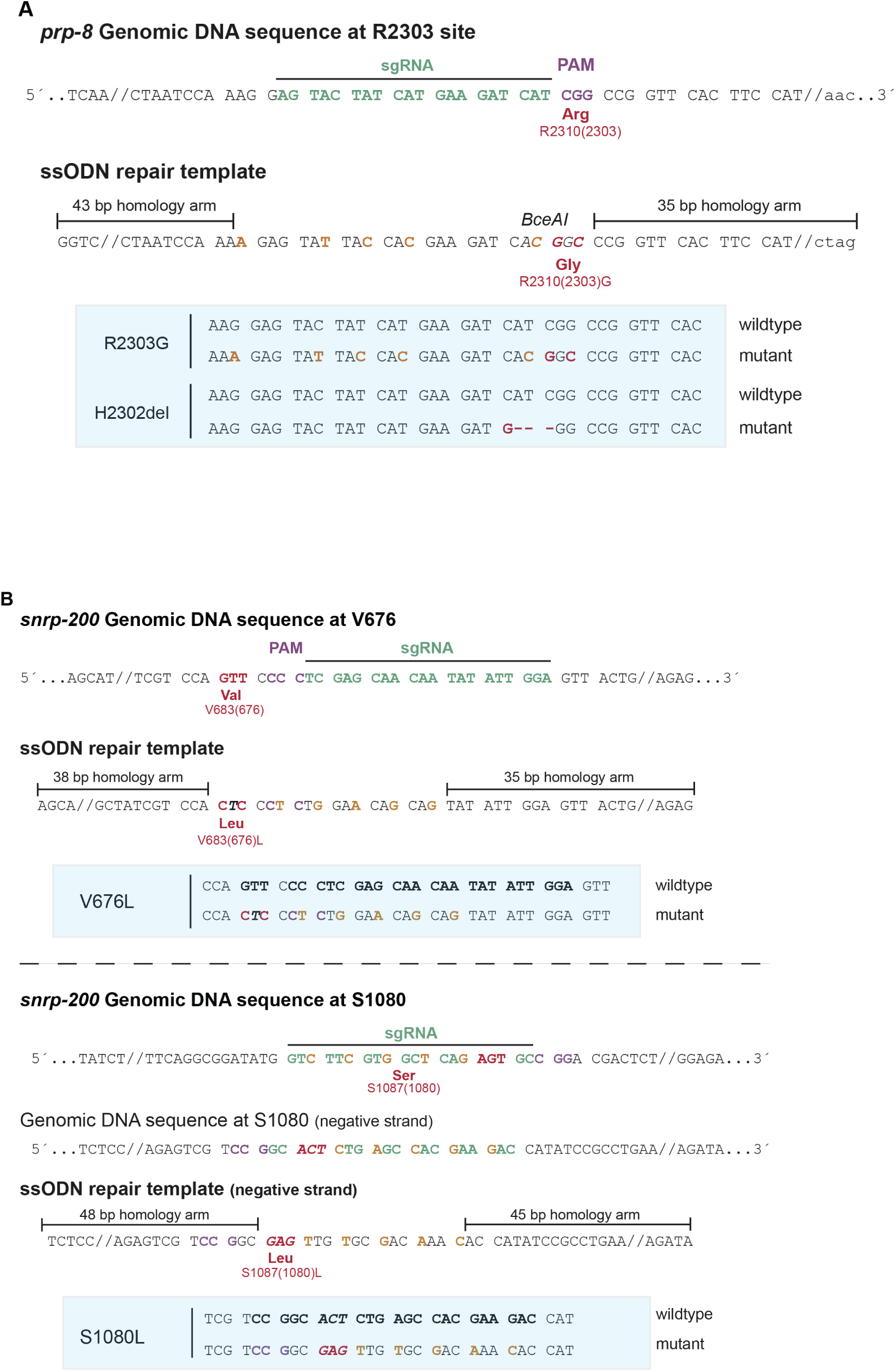
Schematic representation of the CRISPR design to generate s-adRP mutations in *prp-8* and *snrp-200*. PAM sequence (purple), crRNA (green). Silent nucleotide substitutions (orange) were included to prevent re-cutting of Cas9 once the repair template has been incorporated. Nucleotide changes incorporated to produce the mutation are indicated in red.

**Figure S2.**
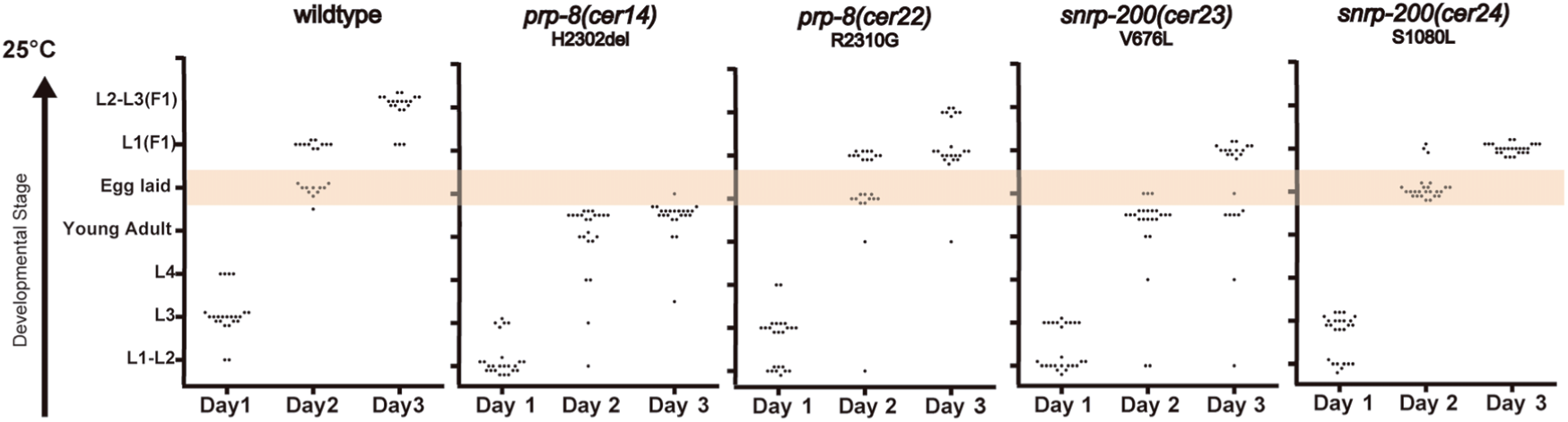
Developmental delay of s-adRP mutants at 20°C. Developmental progress of singled, mutant and wild-type, L1 larvae was monitored every 24 h for 3 days at 20°C. Every dot represents a worm (n=25). Dots inside the shaded area indicate presence of embryos in the plate. Dots inside the green area indicate presence of L1, from the F_1_, in the plate.

**Figure S3.**
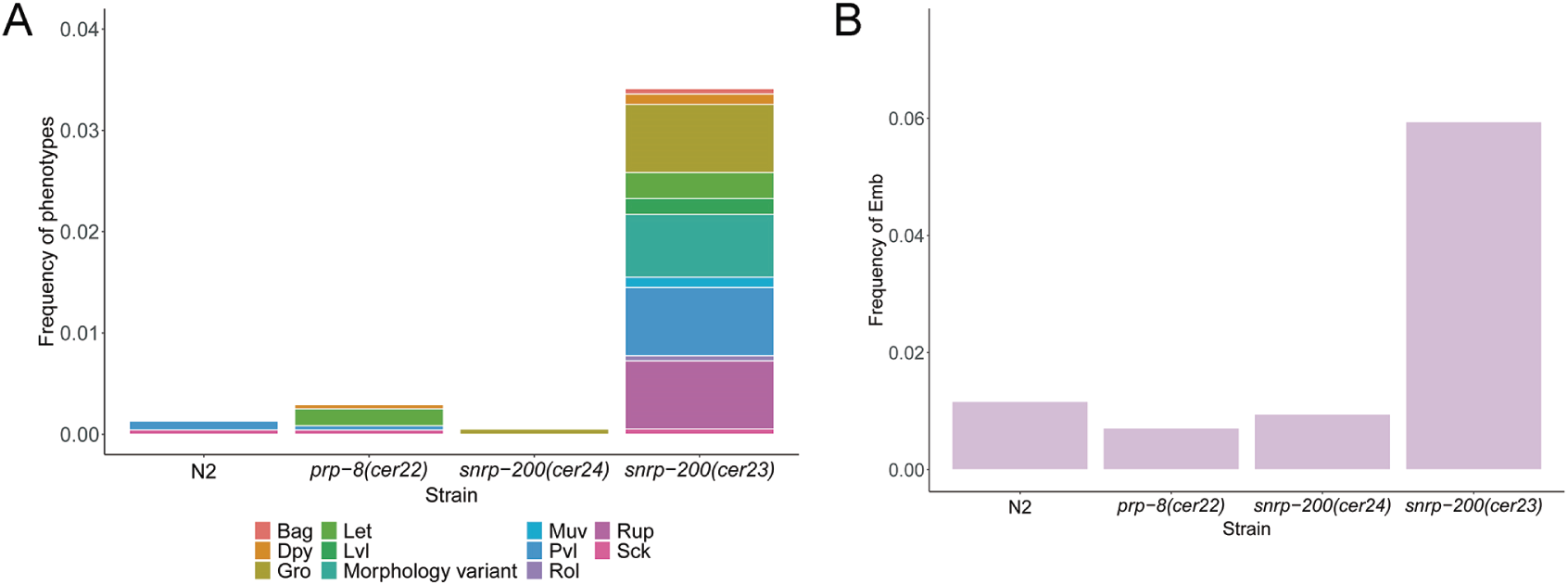
*snrp-200(cer23)* presents higher frequency of Emb and pleiotropic phenotypes at 25°C than *prp-8(cer22)* and *snrp-200(cer24)*. (A) Cumulative frequencies of visually observable phenotypes and (B) frequency of dead embryos of s-adRP mutants (N=1, n≥1753). *prp-8(cer14)* not included as it presents ts-Ste at 25°C. Phenotype abbreviations: Bag: bag of worms, Dpy: dumpy, Gro: growing defects, Let: lethality, Lvl: larval lethality, Muv: multivulva, Pvl: protruding vulva, Rol: roller, Rup: ruptured, Sck: sick.

**Figure S4.**
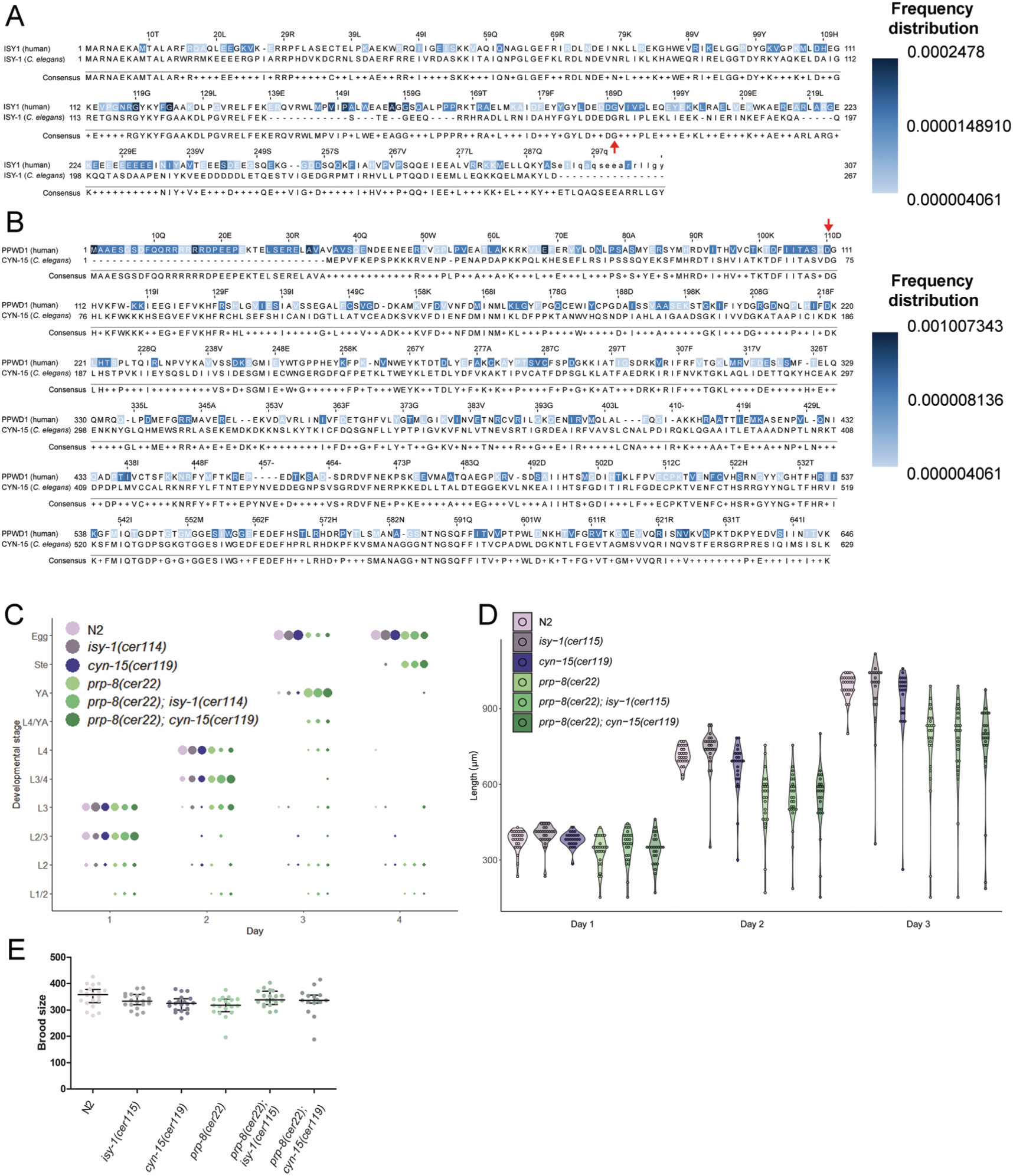
Human polymorphisms of ISY1 (*isy-1* ortholog) and PPWD1 (*cyn-15* ortholog) do not genetically interact with *prp-8(cer22)*. (A) Protein alignment of *C. elegans* ISY-1 with its human ortholog ISY1. (P) Protein alignment of *C. elegans* CYN-15 with its human ortholog PPWD1 (B). Blue boxes mark residues with identified SNPs that cause missense mutations, and the intensity of the color indicates its frequency according to gnomAD (22). Consensus line indicates the conservation of each residue between both organisms. Red arrows indicate residues edited in this study. (C-E) Developmental timing and brood size of the *prp-8(cer22)* with and without (*isy-1(cer114*[G170S]) and *cyn-15(cer119*[D74Q])). (C) Developmental timing. The size of each dot is proportional to the percentage of the population at a given developmental stage starting with a synchronized population and grown at 20°C (N=2, n≥81). (D) Violin plot of the length distribution of a synchronized population over time at 20°C, each dot represents the length of an individual worm (N=1, n≥22). **(E)** Brood size. Each dot represents the brood of an individual worm (N=1, n≥15). Kruskal-Wallis with Dunn’s post-hoc analysis was used with no differences between groups as a result.

**Figure S5.**
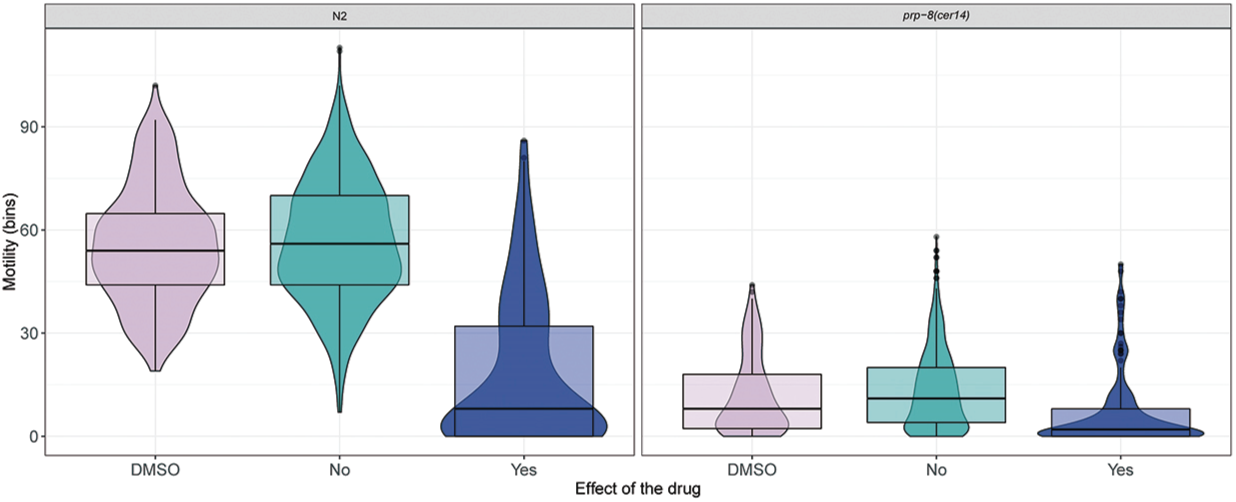
Worms presenting an observable phenotype upon drug treatment have a reduced motility. Violin plot and boxplot of the motility of wild-type (N2) and *prp-8(cer14)* worms, classified by the presence or not of an observable phenotype after drug treatment. *prp-8(cer14*) presents reduced motility even at control conditions. In both, N2 and *prp-8(cer14)*, the motility is drastically reduced in case of drugs causing harmful effects. Note that N2 is not directly comparable to *prp-8(cer14*) due to the contribution of F_1_ in the control worms to the motility, but not in *prp-8(cer14)*. Box-plot indicates the median with the IQR and whiskers ± 1.5 product of IQR. Motility was measured using the wmicrotacker device.

**Figure S6.**
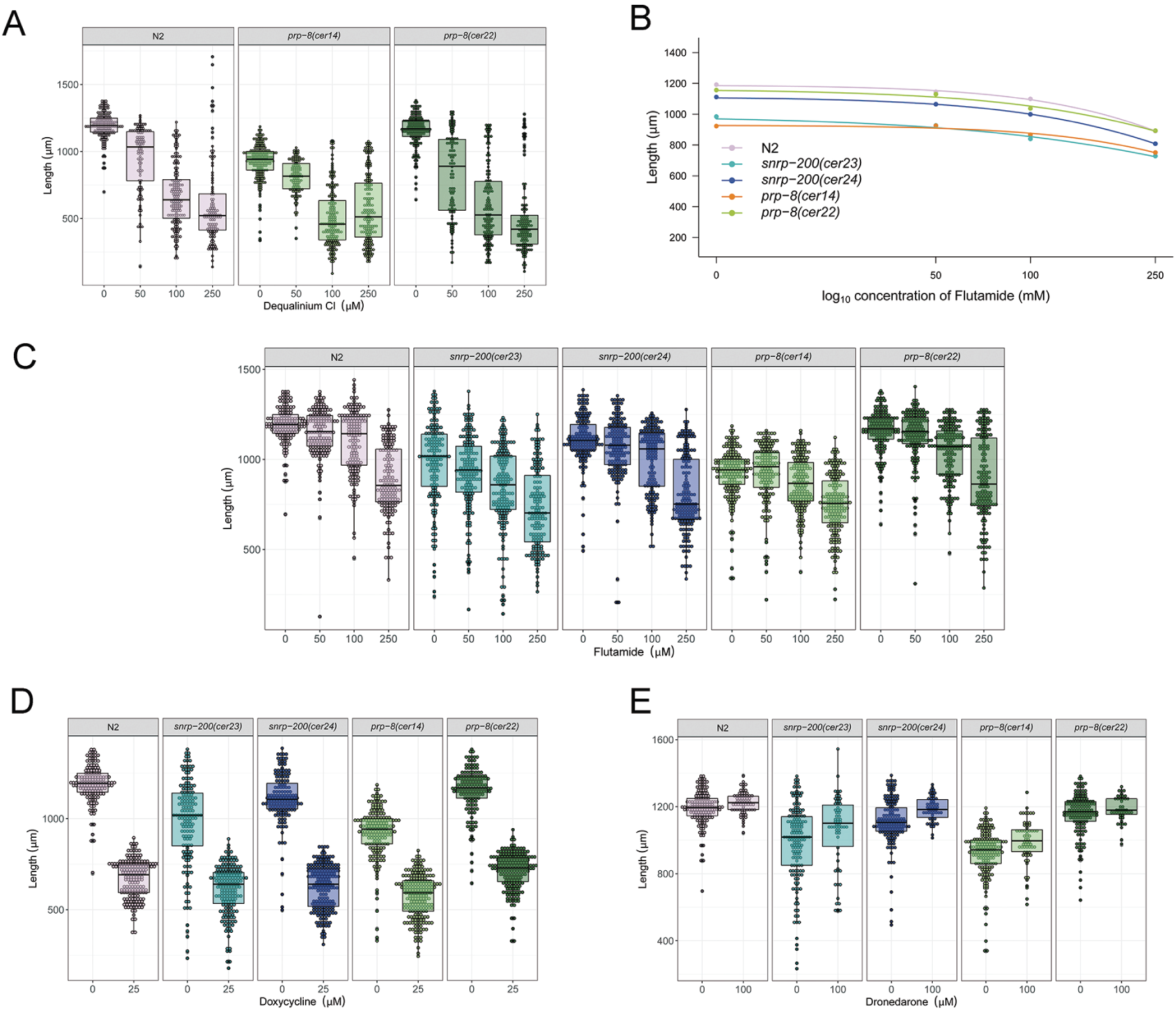
Additional Validation test for candidates from the drug screen. Validation of drugs that induced different response between *prp-8(cer14)* and wild-type (N2) in the primary screen. (A) Worm length upon increasing Dequalinium Chloride concentration in the two *prp-8* s-adRP mutants. Each dot represents the length of an individual worm, box-plot indicates the median with the IQR and whiskers ± 1.5 product of IQR (N=2 at 50 µM, all others N=3, n≥97). Note that *prp-8(cer14)* has a reduced length even at control conditions. (B-C) Worm length measurements upon increasing Flutamide concentration of s-adRP mutants. (B) Each dot represents the median of the length of each condition and the line Log-logistic distribution with lower limit set at 0 (N=3, n≥147). (C) Each dot represents the length of an individual worm, box-plot indicates the median with the IQR and whiskers ± 1.5 product of IQR (N=3, n≥147). (D) Worm length upon Doxycycline hyclate treatment. Each dot represents the length of an individual worm, box-plot indicates the median with the IQR and whiskers ± 1.5 product of IQR (N=3, n≥174). (E) Worm length upon Dronedarone hydrochloride treatment. Each dot represents the length of an individual worm, box-plot indicates the median with the IQR and whiskers ± 1.5 product of IQR (N=1, n≥44).

**Table S1: List of genes tested in the RNAi screen with the phenotypes obtained. (xslx file)**

All validated RNAi clones are shown with each source (Ahringer library (A) or ORFeome based (V)), *C. elegans* locus with its yeast and Human orthologs, in which spliceosome complex it participates, from which class or family of genes it is and the observed phenotype in *cer22* and *cer24* screens. The validated interactors are marked in yellow. Adapted from Serrat. et al 2019, (manuscript in preparation) S4 table. Phenotype abbreviations: Ste: sterility, Rbs: reduced brood size, Gro: developmental delay, Lva: larval arrest, Pvl: protruding vulva, Let: lethality, Lvl: larval lethality, Emb: embryonic lethal, Sck: sick, Rup: ruptured. “p” before a phenotype indicates “partial”.

**Table S2: List of drugs used in the primary screen, phenotypes seen in the primary screen and the re-testing and the motility values.**

**(xslx file)**

Phenotype abbreviations: Ste: sterility, Rbs: reduced brood size, Gro: developmental delay, Lva: larval arrest, Pvl: protruding vulva, Let: lethality, Lvl: larval lethality, Emb: embryonic lethal, Sck: sick, Rup: ruptured, Egl: egg laying defective, Dpy: dumpy. “p” before a phenotype indicates “partial”.

